# Cyclic and pseudo-cyclic electron pathways play antagonistic roles during nitrogen deficiency in *Chlamydomonas reinhardtii*

**DOI:** 10.1101/2023.01.18.524499

**Authors:** Ousmane Dao, Adrien Burlacot, Felix Buchert, Marie Bertrand, Pascaline Auroy, Carolyne Stoffel, Jacob Irby, Michael Hippler, Gilles Peltier, Yonghua Li-Beisson

**Author notes:** **Correspondence to:** Yonghua Li-Beisson or Ousmane Dao. **Competing Interest Statement**: There are no conflicts of interest.

## Abstract

Nitrogen (N) scarcity is a frequently encountered situation that constrains global biomass productivity. In response to N deficiency, cell division stops and photosynthetic electron transfer is downregulated, while carbon storage is enhanced. However, the molecular mechanism downregulating photosynthesis during N deficiency and its relationship with carbon storage are not fully understood. The Proton Gradient Regulator-like 1 (PGRL1) controlling cyclic electron flow (CEF) and Flavodiiron proteins involved in pseudo-(CEF) are major players in the acclimation of photosynthesis. To determine the role of PGRL1 or FLV in photosynthesis under N deficiency, we measured photosynthetic electron transfer, oxygen gas exchange and carbon storage in *Chlamydomonas pgrl1* and *flvB* knockout mutants. Under N deficiency, *pgrl1* maintains higher net photosynthesis and O_2_ photoreduction rates, while *flvB* shows a similar response compared to control strains. Cytochrome *b_6_f* and PSI are maintained at a higher abundance in *pgrl1*. The photosynthetic activity of *flvB* and *pgrl1 flvB* double mutants decreases in response to N deficiency similar to the control strains. Furthermore, the preservation of photosynthetic activity in *pgrl1* is accompanied by an increased accumulation of triacylglycerol depending on the genetic background. Taken together, our results suggest that in the absence of PGRL1-controlled CEF, FLV-mediated PCEF maintains net photosynthesis at a high level and that CEF and PCEF play antagonistic roles during N deficiency. It further illustrates how nutrient status and genetic makeup of a strain can affect the regulation of photosynthetic energy conversion in relation to carbon storage and provides new strategies for improving lipid productivity in algae.

**Significance statement:** Nitrogen (N) deficiency, an often-encountered phenomenon in nature, results in growth arrest, downregulation of photosynthesis and massive carbon storage in microalgae. However, more mechanistic insights involved in tuning photosynthetic electron transfer during N deficiency are required. Here, we provide evidence that a well-conserved protein in chlorophytes, the Proton Gradient Regulator-like 1 (PGRL1), is a key regulator of photosynthesis during N deficiency. In its absence, cells exhibited sustained photosynthesis thanks to the Flavodiiron (FLV) proteins. We propose that both PGRL1 and FLV, by having antagonistic roles in N deficiency, manage the redox landscape, carbon storage and biomass production. Our work revolves around the current paradigm of photosynthesis regulation during N deficiency and provides a new framework for improving biomass production and carbon storage in microalgae for biotechnological purposes.

## Introduction

Nitrogen (N) deficiency is one of the most harsh environmental situation that constrains global primary biomass productivity in all ecosystems (1–3). Under N deficiency, cell division stops and photosynthetic CO_2_ assimilation is downregulated, the carbon and energy being used to synthesize starch and triacylglycerols (TAGs) (4–7). The downregulation of photosynthetic electron transfer (PET) reactions, together with a re-routing of the excess reducing power towards carbon storage, prevents the over-production of reactive oxygen species, thus ensuring cell fitness (8–12).

Due to the variability in their natural habitat, microalgae must constantly adjust the photosynthetic conversion of energy to match the metabolic demand and have therefore developed a set of regulatory mechanisms to fine-tune electron transfer reactions. During photosynthesis, energy in the form of ATP and NADPH is mostly produced by the linear electron flow (LEF) (13). The balance of ATP and NADPH is essential for optimal CO_2_ capture and metabolism to which cyclic electron flow (CEF) and pseudo-cyclic electron flow (PCEF) play a critical role (14–18). Two pathways of CEF around PSI have been described in the green microalga *Chlamydomonas reinhardtii* (*Chlamydomonas* hereafter), one involving the type II NADPH dehydrogenase (NDA2) (19), and the other being controlled by the proton gradient regulator 5 (PGR5)/PGR-like 1 (PGRL1) proteins (20).

Recent work in *Chlamydomonas* provided evidence that PGR5 and PGRL1 are involved in CEF (20–22). Hereby PGR5 is required for efficient stromal electron intake into the cytochrome *b*_6_*f* complex (Cyt *b*_6_*f*) (23). Its deletion thereby strongly disturbs the Mitchellian Q cycle (23). Also, deletion of PGRL1 impacts CEF in *Chlamydomonas* (20, 21). Despite the fact that PGRL1 has been implicated in plastoquinone reduction during CEF (24), it appears that PGRL1 is rather important for PGR5 expression and protein stability, as in the absence of PGRL1, PGR5 is strongly diminished, mimicking PGR5 dependent phenotypes (21, 25, 26).

Both CEF pathways reduce the plastoquinone (PQ) pool by using either NADPH or ferredoxin (Fd) as electron donor respectively (19, 24, 27). CEF has been shown to contribute to the acidification of thylakoid lumen by generating an extra proton motive force (*pmf*) in addition to the one produced by LEF (19, 20, 28, 29). PCEF mediated by Flavodiiron proteins (FLVs), by transferring electrons toward O_2_ at the acceptor side of PSI, also contributes to the establishment of the *pmf* and therefore to the lumen acidification (30–32). The *pmf* is used to either (1) produce ATP (33), (2) trigger light energy quenching via a low lumenal pH (34–36), (3) convert HCO_3_ ^-^ into CO_2_ thanks to carbonic anhydrase in the lumen (32) or (4) repress electron transfer at the level the Cyt *b*_6_*f* complex through the photosynthetic control triggered by the low lumenal pH (28, 37–40). Both CEF and PCEF have been shown to be critical under various conditions of light, CO_2_ availability or sulfur deficiency by playing a synergistic role (14, 31, 32, 41, 42).

Despite the importance of CEF and PCEF in response to dynamic environments, little is known about their role during N deficiency. The role of NDA2- involved CEF have been recently addressed during CO_2_-limiting photoautotrophic N deficiency and the knockdown of NDA2 was shown to impair the establishment of non-photochemical quenching (29). The role of PGRL1/PGR5-controlled CEF has been investigated under mixotrophic N deficiency and the lack of PGRL1 was shown to decrease the rate of CEF and TAG production (43). However, the role of PGRL1 during photoautotrophic N deficiency and the possible bioenergetic interactions between carbon/energy sinks (TAG and starch) and CEF or PCEF pathways that generate ATP has not yet been explored so far. Considering that massive carbon reallocation occurs under N deficiency (4), the bioenergetics governing this reallocation are likely critical. Fully understanding photosynthesis regulatory pathways and how they affect biomass production and reserve formation is needed towards engineering photosynthesis and carbon storage in conditions of nutrient deficiency.

Here, we evaluated the contribution of PGRL1/PGR5-controlled CEF in the regulation of photosynthesis during N deficiency and evaluate its metabolic consequences on carbon storage in *Chlamydomonas* cells grown in photoautotrophic conditions under non-limiting CO_2_ concentrations (using 1% CO_2_-enriched air), conditions which favor the accumulation of carbon reserves (7, 44, 45). By monitoring photosynthetic activity based on chlorophyll fluorescence and O_2_ exchange rate measurements, we observed high net photosynthetic activity in PGRL1-deficient strains under N deficiency compared to the control strains. Furthermore, the lack of PGRL1 in 137AH genetic background resulted in an over-accumulation of TAGs. Those effects being suppressed in double mutants deficient in both PGRL1 and FLVB. We conclude that FLVs, by maintaining a strong PCEF activity in the *pgrl1* mutants, catalyze high photosynthetic rates. Finally, we discuss how modulating photosynthetic electron flow could constitute an efficient strategy to improve photosynthesis and eventually boost further TAG production under N deficiency.

## Results

### Photosynthetic activity sustained longer in *pgrl1* under N deficiency

To assess the role of the PGRL1/PGR5-controlled CEF in the regulation of photosynthesis during photoautotrophic N deficiency, we simultaneously monitored chlorophyll fluorescence and O_2_ exchange following a dark-light-dark transition in N-replete and N-deprived cells. We compared the photosynthesis efficiency of a PGRL1-deficient strain (*pgrl1_137AH_)* with its control wild-type strain (137AH) and a complemented line (20) (**Fig. 1 and Supplemental Fig. S1 and S2**). Under N replete conditions, both chlorophyll fluorescence and O_2_ exchange patterns were mostly similar in *pgrl1_137AH_* and control cells (**Fig. 1*A, C*, *E* *and F* and Supplemental Fig. S1 *B-C***). After 2 days of N deficiency, the PSII operating yield measured in the light decreased by about 75% in the wild-type and only by 35% in *pgrl1* (**Fig. 1A, B, and *E*; Supplemental Fig. S1*C* and *E***). Conversely, *pgrl1_137AH_* showed twice higher net O_2_ evolution and light-dependent O_2_ uptake as compared to the wild-type (**Fig. 1*D*, *F***). As previously reported (5, 46), the dark O_2_ consumption rate was stimulated in the 137AH strain during N deficiency, this effect being reduced in *pgrl1_137AH_* **(Fig. 1*D*; Supplemental Fig. S2*B***). By using *pgrl1_137AH_*complemented line *pgrl1::PGRL1*-2 (20), we noticed a full recovery of all the observed phenotypes (**Fig. 1 *E*-*G;* Supplemental Fig. S1 and S2**). To further strengthen the phenotype-genotype relationship, we generated an additional *pgrl1_CC125_* mutant in the CC125 wild-type genetic background using CRISPR-Cas9. In line with our observation in the 137AH background **(Fig. 1*E-F*)**, we detected a sustained retention of photosynthetic PSII yield and higher net O_2_ evolution in the *pgrl1_CC125_* during N deficiency compared to CC125 (**Supplemental Fig. S3**). The higher O_2_ consumption under N deficiency in the dark-adapted 137AH strain from **Fig. 1*D*** was independent of PGRL1 in the CC125 background (**Supplemental Fig. S3 *D-E***). The preservation of photosynthetic activity in *pgrl1* suggests that PGRL1-controlled CEF contributes to repressing the PET reactions under photoautotrophic N deficiency.

**Figure 1.**
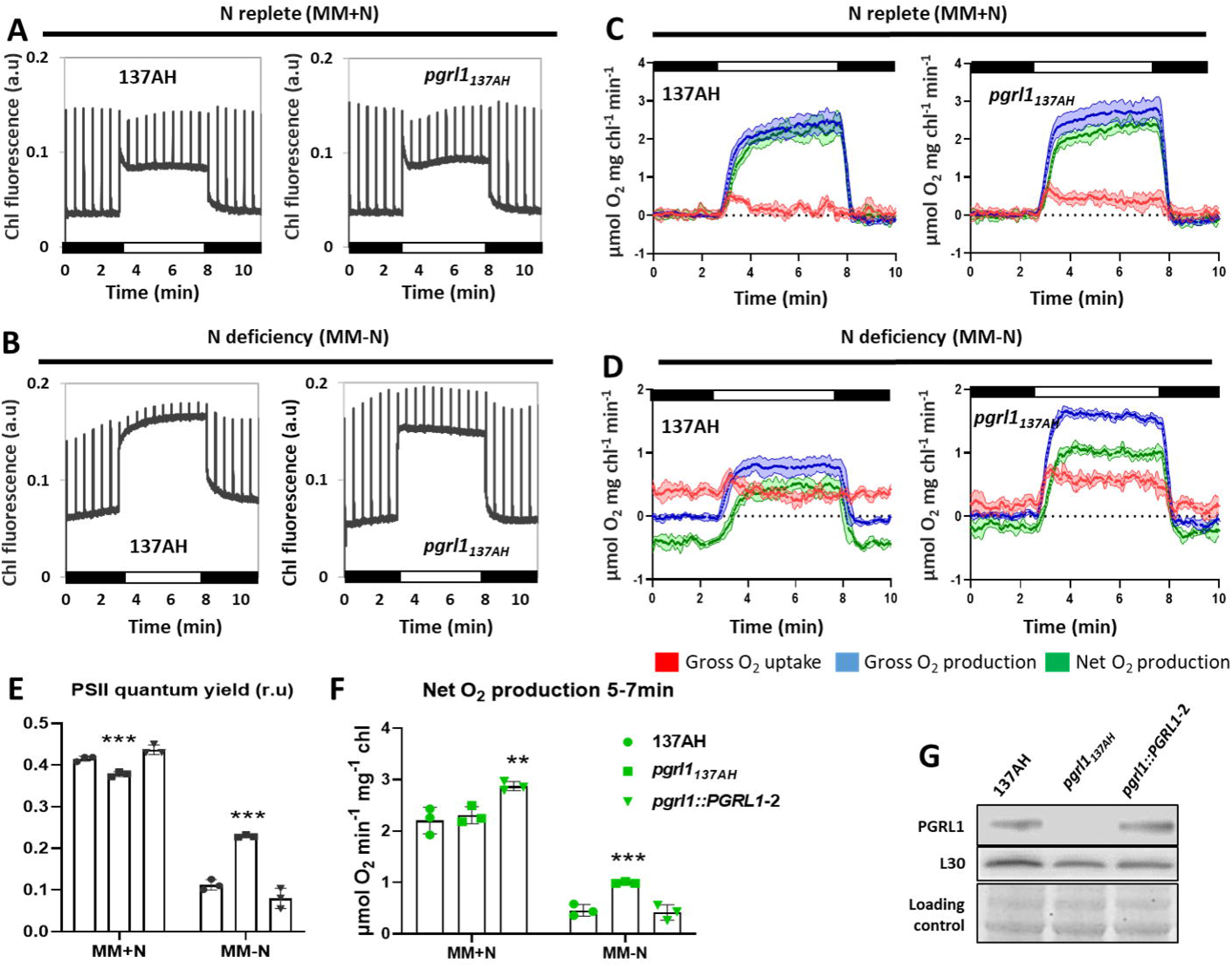
*pgrl1* showed sustained photosynthesis after 2 d of N deficiency. **A-B**, Chlorophyll fluorescence was measured using a Dual-PAM during the dark- light-dark transition from N-replete (**A**) and N-deficient (**B**) conditions. **C-D**, O_2_ exchange rates were measured using a MIMS during the dark-light-dark transition from N-replete (**C**) and N-deficient (**D**) conditions. Net O_2_ evolution (green) was calculated as gross O_2_ evolution (blue) – gross O_2_ uptake (red). **E**, PSII quantum yield before and after 2 d of N deficiency measured using green actinic light (1250 µmol photon m^-2^ s ^-1^, green LEDs) and calculated as ΦPSII= (F_M_’- Fs)/ F_M_’ with F_M_’ the fluorescence value after saturating pulse, Fs the stationary fluorescence during actinic light exposure. **F**, The Net O_2_ evolution before and after 2 d of N deficiency in the wild-type 137AH, *pgrl1_137AH_* and the complemented line *pgrl1::PGRL1-2*. **G**, Immunoblot analysis of PGRL1 accumulation in the wild-type 137AH, *pgrl1* and the complemented line. L30 antibody was used as a positive control. Coomassie blue staining was used as the loading control. Protein samples were obtained from 2 d N- starved cells. N-replete wild-type 137AH, *pgrl1_137AH_,* and *pgrl1::PGRL1-2* cells cultivated photoautotrophically with 1% CO_2_ in air under continuous light (50 μmol photons m^−2^ s^−1^) were transferred into N-free media for 2 d prior to measurements. Shown are an average of at least three biological replicates ± SD. Asterisks represent statistically significant differences compared to the control 137AH (* *p*<0.05, ** *p*<0.01 and *** *p*<0.001) using one-way ANOVA.

### The PGRL1 modulates the PSI donor and acceptor side during N deficiency

The PGRL1/PGR5-controlled CEF is a determinant of the downstream PSI electron fate which prompted us to monitor P700, the primary PSI donor (**Fig. 2; Supplemental Fig. S4**). P700 forms three populations and only the first two are photo-oxidizable: 1. PSI donor shortage generates P700^+^ in the environmental light, 2. PSI yield quantifies P700^-^ during environmental acclimation that converts within the ms range to P700^+^ upon a saturating light pulse, and 3. PSI acceptor side limitation represents a redox-inactive P700 pool (see Materials and Methods). In the absence of PGRL1/PGR5-controlled CEF the redox-inactive P700 pool is enhanced under CO_2_-limiting conditions (47). This was also observed in the 137AH genetic background when *pgrl1* was grown photoautotrophically under 1% CO_2_ (**Fig. 2*A, B***). The failure to produce P700^+^ in the light gradually developed within 0.5 s and persisted throughout the illumination period (**Fig. 2*A, B***). However, *pgrl1* did not display lower PSI yields, typically detected in ambient CO_2_ (48), since the diminished P700^+^ was due to lower donor side limitation (**Fig. 2*B***). The P700 differences in *pgrl1* relied on PSII activity, pointing to a LEF/CEF entanglement. N depletion further increased the P700^+^ pool in a strictly donor side-dependent fashion in both wild-type and mutant. Nevertheless, *pgrl1* displayed twice the PSI acceptor side limitation when PSII was active. Moreover, the mutant showed a delayed P700^+^ formation within the first 200 ms of light when PSII was inhibited (**Fig. 2*A***). The latter feature was also displayed in N-deficient CC125 under DCMU conditions (**Supplemental Fig. S4*A***). This genetic background showed identical photo-oxidizable P700 pools between wild-type and mutant in the presence of N, with *pgrl1* producing slightly higher acceptor side limitation (**Supplemental Fig. S4*B***). N depletion came at the expense of PSI yields and *pgrl1* failed to alleviate the PSI acceptor side through sufficient donor side induction. These results suggest that PGRL1-controlled CEF facilitates the onset of photosynthetic control during photoautotrophic N deficiency by inducing a donor side limitation at PSI to alleviate its acceptor side redox pressure.

**Figure 2.**
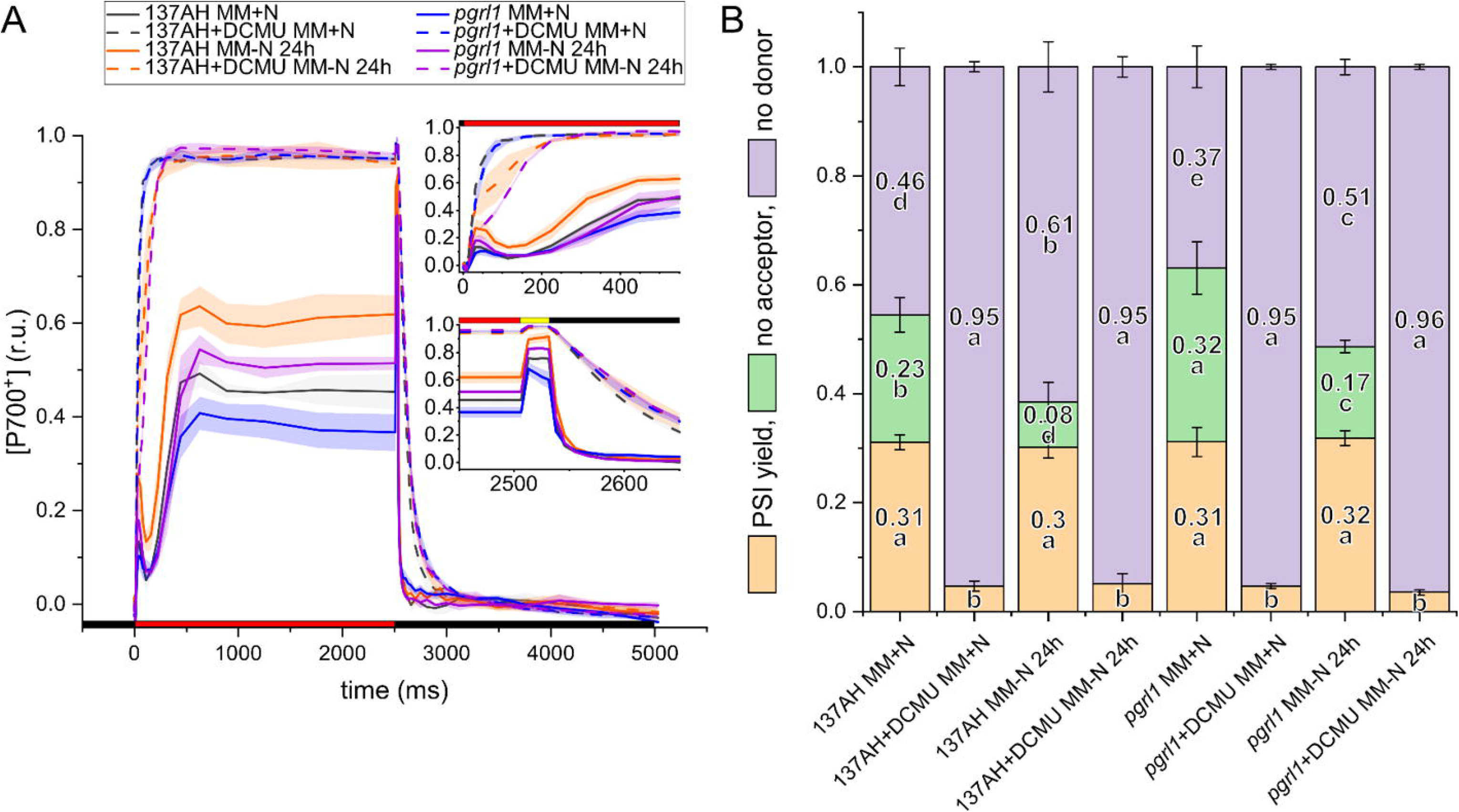
PSI oxidation is facilitated by PGRL1 under N deficiency. The influence of PGRL1 on the redox state of the primary PSI donor, P700, was monitored in the genetic background 137AH **(A-B)** and CC125 **(Supplemental Fig. S4)**. Panel A shows raw kinetics (red/yellow/black bars: 490/3000/0 μmol photons m^−2^ s^−1^) in the presence and absence of PSII inhibitor DCMU (means ± SD of 4 biological replicates). Panel B quantifies the corresponding P700 pools at the end of the 2.5-s light period (letters indicate parameter-specific significances using one-way ANOVA/Fisher-LSD, *p*<0.05). At the time of quantification, P700 remained photo-oxidizable by a saturating pulse (PSI yield), was redox inactive (no acceptors) or was pre-oxidized (no donors). N-replete cells of wild-type 137AH and *pgrl1_137AH_*were cultivated photoautotrophically with 1% CO_2_ in air under continuous light (50 μmol photons m^−2^ s^−1^), and then transferred into N-free media for 24 h prior to measurements.

### FLV-mediated O_2_ photoreduction drives photosynthesis in *pgrl1* under N deficiency

Because light-dependent O_2_ uptake can lead to the generation of extra ATP and support CO_2_ fixation but at various levels depending on the mechanism involved (18), we sought to test the nature of the light-dependent O_2_ uptake mechanism activated in *pgrl1* mutants during N deficiency (**Fig. 1*D***). FLVs have been reported to be a major O_2_ uptake mechanism in the light (31, 32, 49). *pgrl1 flvB* double mutants impaired in both PGRL1 and FLVB have been recently generated by crossing the *pgrl1_137AH_* with a FLVB deficient strain (*flvB21*) as described in (32). Therefore, we used the parental lines as control strains (*flvB-21* and *pgrl1_137AH_*) as well as sibling strains from the progeny of the crossing (WT1 and WT3) harboring both FLV and PGRL1 proteins (**Supplemental Fig. S1A)**. The two mutants (*pgrl1 flvB-3* and *pgrl1 flvB-5)* and control sibling (WT1 and WT3) strains used throughout the manuscript have been chosen for having similar maximal photosynthesis under N-replete conditions (V_Max_) (18, 32). For all the experiments, we kept the *pgrl1_137AH_*alongside with its control 137AH and *pgrl1::PGRL1*-2; *flvB* mutants (*flvB21*, *flvB208* and *flvB308*) alongside with their control CC4533 and finally *pgrl1 flvB-3* and *pgrl1 flvB-5* alongside WT1 and WT3 as control. We evaluated steady-state O_2_ exchange rate in *pgrl1_137AH_*, *flvB* mutants as well as in the *pgrl1 flvB* double mutants after 2 days of N deficiency (**Fig. 3**). The light-dependent O_2_ uptake was highly increased in *pgrl1_137AH_* but strongly impaired in *flvB* and *pgrl1 flvB* mutants (**Fig. 3A; Supplemental Fig. S2**) which mirrors what has been reported for N-replete conditions in the presence of atmospheric CO_2_ level (31, 32). Interestingly, while the gross O_2_ uptake, the gross O_2_ and the net O_2_ evolution measured during steady-state photosynthesis remained high in *pgrl1_137AH_*, no difference was detected between *flvB*, *pgrl1 flvB* mutants and their respective control lines (**Figure 3*B-D* and Supplemental Fig. S2).** Taken together, these data show that the absence of FLV-mediated O_2_ photoreduction (in *pgrl1 flvB* mutants) suppresses the effect of N deficiency observed in *pgrl1_137AH_* strain. We conclude that FLV-mediated O_2_ photoreduction, by maintaining PET reactions in *pgrl1,* allows net photosynthesis to be maintained at a high-level during N deficiency.

**Figure 3.**
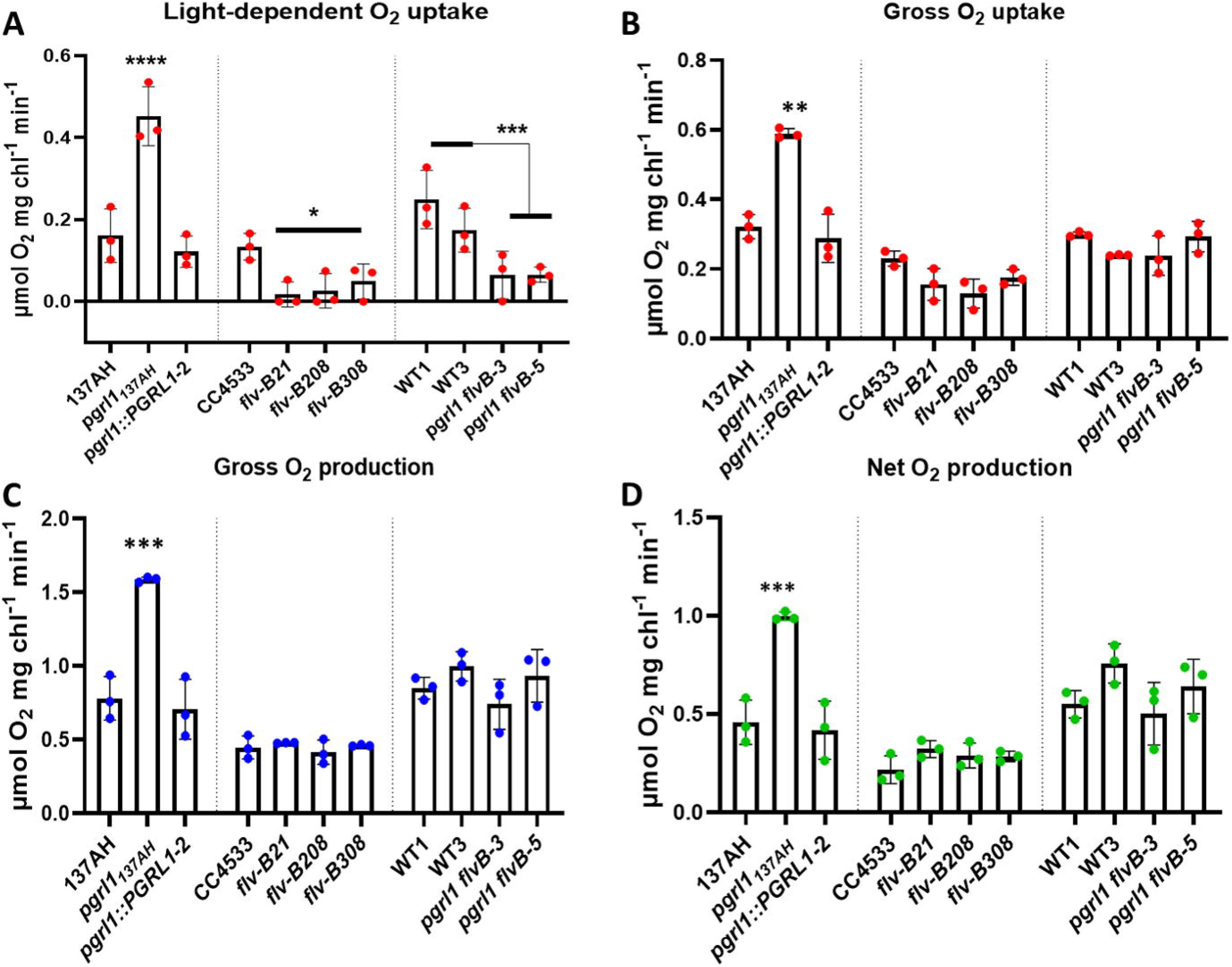
The photosynthesis in *pgrl1* is driven by FLV-mediated O_2_ photoreduction under N deficiency. **A,** Light-dependent O_2_ uptake attributed to FLV activity and calculated as the difference between the O_2_ uptake during the 1^st^ min of illumination and dark O_2_ uptake as shown by the **Supplemental** Figure 2. **B**, Gross O_2_ uptake measured between 5-7min as shown by the **Supplemental** Figure 2. **C,** Gross O_2_ evolution measured between 5-7min as shown by the **Supplemental** Figure 2. **D**, Net O_2_ evolution measured between 5-7min as shown by the **Supplemental** Figure 2. O_2_ exchange rates were measured using a MIMS in the presence of [^18^O]-enriched O_2_. Net O_2_ evolution (green) was calculated as gross O_2_ evolution (blue) – gross O_2_ uptake (red). Total proteins were extracted from N-replete and 2 days N-deprived cells and samples were loaded at equal total proteins amounts as shown on Coomassie blue staining. Cells cultivated photoautotrophically with 1% CO_2_ in air under continuous light of 50 μmol photons m^−2^ s^−1^ were transferred into N-free media for 2 days prior to measurements or sampling for immunoblot. Shown are an average of at least three biological replicates ± SD. Asterisks represent statistically significant difference compared to the wild-type strains (* *p*<0.05, ** *p*<0.01, *** *p*<0.001 and **** *p*<0.0001) using one-way ANOVA.

### Cyt *b**_6_**f and* PSI subunits accumulate to higher levels in *pgrl1* under N deficiency

To gain insights into the mechanisms behind the high photosynthetic activity in *pgrl1* during N deficiency, we compared the relative abundance of representative catalytic core subunits from photosynthetic complexes in the *pgrl1_137AH_* and the 137AH wild-type during N deficiency (**Fig. 4*A and D***). Additionally, we also investigated potential changes in mitochondrial electron transport chain by probing Complex II (Cox IIB). We observed a sustained retention of the PSI subunit PsaD and Cyt *f* subunit of Cyt *b_6_f* in *pgrl1_137AH_* compared to the 137AH (**Fig. 4*A and D***). Higher amounts of PsaD and Cyt *f* were observed in *pgrl1_137AH_* under N replete without any functional effect on PSII yield and Net O_2_ evolution (**Fig. 1*E-F*; 4*A and D*)**. As for the stimulated dark O_2_ consumption **(Fig. 1*D*)**, the amount of mitochondrial respiratory chains component Cox IIB was increased in the 137AH whereas it remains stable in the *pgrl1_137AH_* during N deficiency (**Fig. 4*A***). It is worth noting that the hallmark of autophagy, i.e., ATG8 was barely detectable in the *pgrl1_137AH_*whereas it was highly induced in the 137AH under N deficiency (**Fig. 4*A, D***). All the other proteins tested accumulated to a similar amount in *pgrl1_137AH_* and 137AH (**Fig. 4*A***). Note that higher amounts of Cyt *f* were also observed in CRISPR-generated *pgrl1_CC125_* mutant as compared to CC125 in response to N deficiency (**Supplemental Fig. S3*E*)**. We conclude from these experiments during N deficiency that the higher photosynthetic activity observed in mutants impaired in PGRL1 might result from a lower decrease in the amounts of PSI and Cyt *b_6_f* complexes.

**Figure 4.**
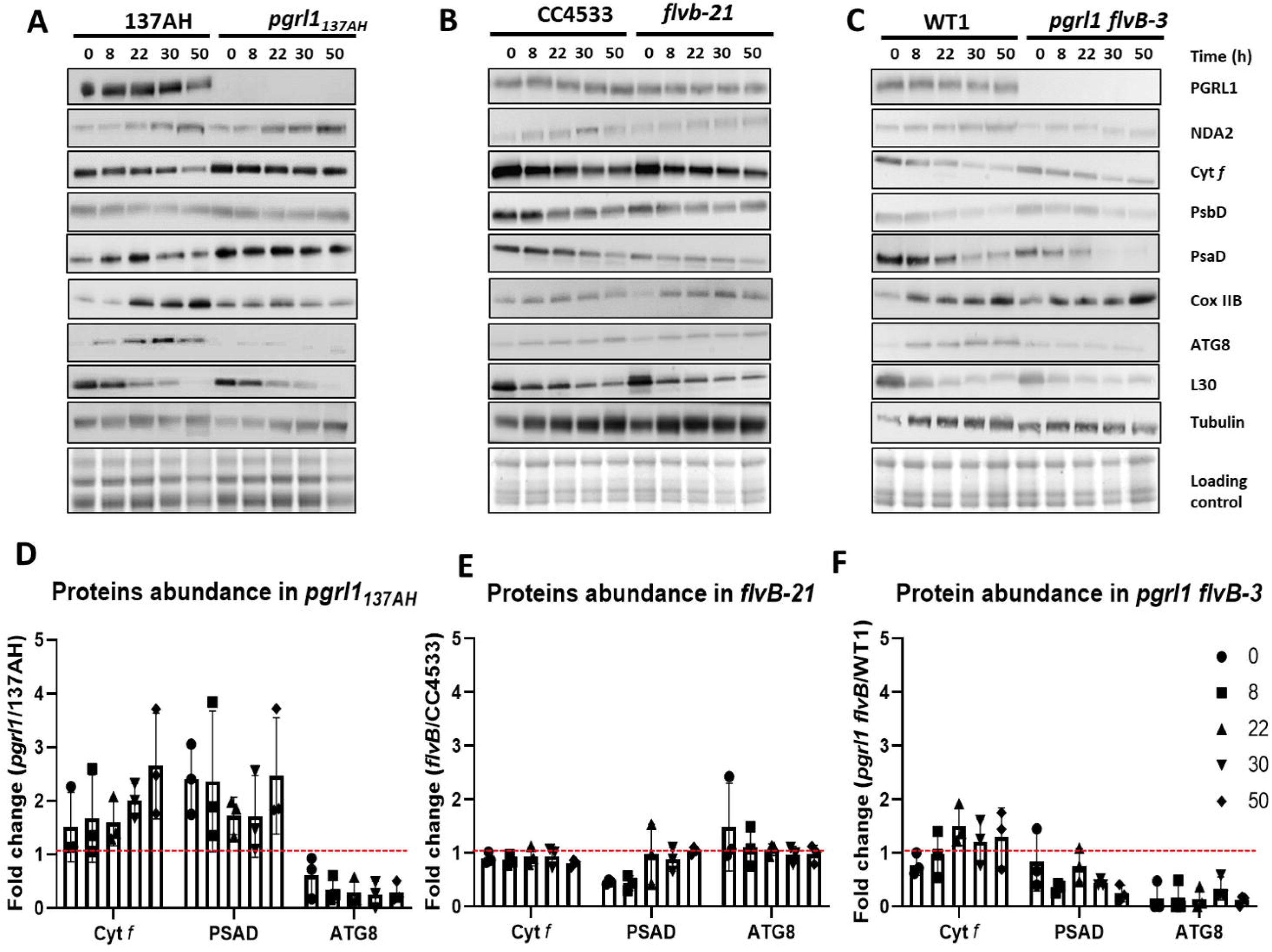
*pgrl1* accumulated more cytochrome Cyt *b_6_f* and PsaD during N deficiency. **A-C**, Immunodetection of photosynthetic proteins in the *pgrl1_137AH_*, *flvB-21* and *pgrl1 flvB-3* as compared to their controls during N deficiency. Cells were harvested at 0, 8, 22, 30 and 50 hours of N deficiency. Samples were loaded at equal total protein amounts as shown by Coomassie blue staining. Shown are representative images of three biological replicates. The α-tubulin was used as housekeeping protein whereas ATG8 and the chloroplast 50S ribosomal large subunit L30 are controls of N depletion (81, 82). **D-F**, Histogrammes showing the abundance Cyt *f*, PsaD and ATG8 proteins in the *pgrl1_137AH_*, *flvB-21* and *pgrl1 flvB-3*during N deficiency in the wild-type cells. Shown represent the ratio of mutants over control strains. Data are average of at least three biological replicates.

To test whether the retention of the PSI and Cyt *b_6_f* observed in the *pgrl1_137AH_* are consequences of the existence of a FLV activity, similar immunoblot analyses were performed in the *flvB-21* and *pgrl1 flvB-3* mutants. Consistent to the photosynthetic activity, immunoblot analyses showed a similar accumulation of tested proteins in the *flvB-21* and *prgl1 flvB-3* compared to controls with the exception of lower PsaD and ATG8 in *prgl1 flvB-3* (**Fig. 4 *B-C* and *E-F***).

### Defects in CEF and PCEF altered carbon storage during N deficiency

Emerging literature suggests that alterations in energy management pathways affect not only biomass production but also its composition (16, 17, 44, 50). Starch and TAGs are major forms of carbon storage in *Chlamydomonas* during N deficiency. Since the *pgrl1_137AH_* mutant shows a stronger net photosynthetic rate, we measured its ability to accumulate storage compounds. Starch accumulation was similar in *pgrl1_137AH_* and control lines under both N replete and deficiency (**Fig. 5*A*; Supplemental Fig. S5*A* and *S6A***). In contrast, twice more TAG accumulation was observed in *pgrl1_137AH_* compared to the control lines under N deficiency whereas no difference was observed under N-replete condition (**Fig. 5*D*; Supplemental Fig. S5*B* and S6*D***). Lipid droplet imaging confirmed the accumulation of higher TAG amounts (**Supplemental Fig. S5*D***). Intriguingly, the *PGRL1* knockout mutant did not induce TAGs or starch accumulation in response to N deficiency in the CC125 genetic background (**Supplemental Fig. S6 *G-J***), which may be related to the fact that photosynthesis preservation (PSII yield and net O_2_ evolution) was lower in the *pgrl1_CC125_* background as in the *pgrl1_137AH_* background (**Supplemental Fig. S3**) thus limiting reserve accumulation, or indicating the existence of a metabolic limitation for storage compound accumulation in the CC125 background.

**Figure 5.**
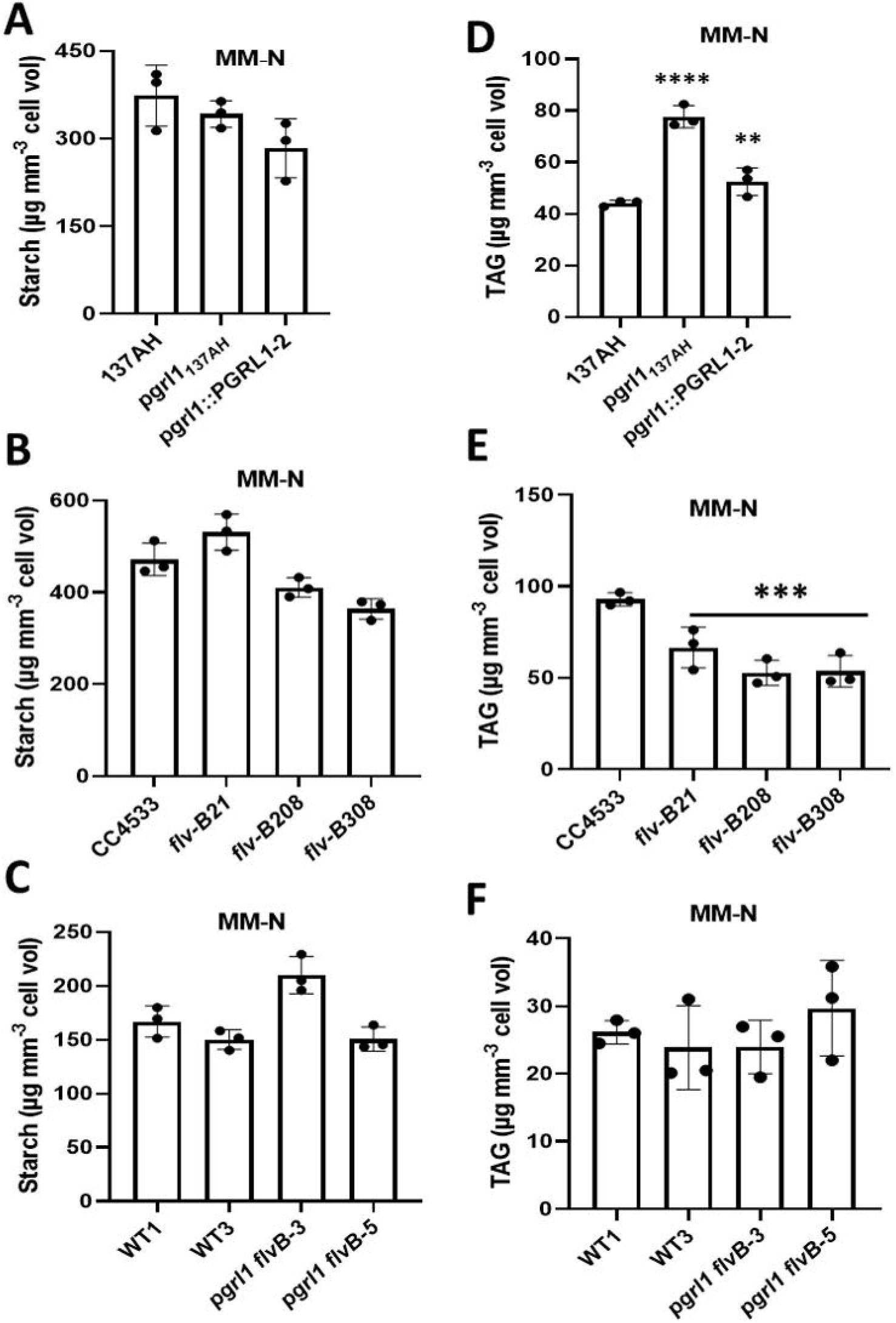
*pgrl1* over accumulated TAGs under N deficiency. Starch **(A-C)** and TAG (**D-F**) quantification in N-deprived cells of *pgrl1_137AH_*, *flvB* and *pgrl1 flvB* mutants respectively. Cells were cultivated photoautotrophically with 1% CO_2_ in air under continuous light of 50 μmol photons m^−2^ s^−1^. For N deficiency, cells were transferred into N-free media for 2 d prior to sampling for starch and TAG. Shown are an average of at least three biological replicates ± SD. Asterisks represent statistically significant difference comparing mutants with their control strains (* *p*<0.05, ** *p*<0.01 and *** *p*<0.001) using one-way ANOVA.

Under N replete, the *flvB* single mutants accumulated higher amounts of starch than the CC4533 wild-type (**Supplemental Fig. S6*B***) but accumulated lower amounts of TAG under N deficiency (**Fig. 5*E***). The double mutants *pgrl1_137AH_ flvB* accumulated similar amounts of TAG and starch as compared to control strains either in N replete or N deprived conditions (**Fig. 5*C, F*; Supplemental Fig. S6*C, F***).

We then evaluated to which extent observed changes in photosynthesis affect biomass accumulation, by following changes in cellular volume as a proxy for biomass accumulation. After 6 days of N deficiency, the cell volume increased twice as fast in the *pgrl1_137AH_* compared to the control lines (**Supplemental Fig. S5*C***). We conclude from the above experiments that the higher photosynthetic activity in *pgrl1_137AH_* results in increased biomass and TAG productivity under N deficiency.

Altogether, our data suggested that during phototrophic N deficiency, photosynthetic electron flow mediated by FLVs could trigger higher TAG accumulation while PGRL1 tends to constrain it, likely via a photosynthetic control-related process or specific protease pathways that target Cyt *b*_6_*f*. We propose that both PGRL1 (CEF) and FLVs (PCEF), by having an antagonistic role during N deficiency, manage the redox landscape, carbon storage and biomass production. It is worth pointing out that this modification of metabolism as a consequence of redox management depends not only on genetic background (137AH versus CC125) but also on the environmental state (+N versus –N).

## Discussion

The ability of microalgae to coordinate their energy conversion (from light to chemical energy) to meet the metabolic demand is crucial for their survival in a constantly fluctuating environment. Mechanisms involved in photosynthesis regulations have been abundantly studied in response to light or CO_2_ levels, but not much is known during nutrient deficiency when a massive re-orientation of metabolic pathways occurs. In the absence of N, the major cellular energy sinks (cell division, protein biosynthesis and photosynthetic CO_2_ fixation) are restricted whereas the carbon and energy are stored as TAGs or starch. Moreover, the PET reactions are downregulated under N deficiency. However, we do not know whether the downregulation of PET is a mean to match cells energy status with the limited metabolic demand. In other words, how the metabolism accommodates when photosynthesis remains high during N deficiency is not well understood. In this study, we demonstrated that PGRL1, which controls a CEF pathway, tunes PET during N deficiency. This tuning favors the photosynthetic control mechanism via Cyt *b*_6_*f* to limit PSI donor availability (see below for details) and might affect energy-transducing enzyme ratios at least in chloroplasts. Lack of PGRL1 resulted in sustained photosynthetic activity up to 2 days of N deficiency. The misregulation of the PGRL1/PGR5-controlled CEF is notorious for decreasing the photosynthetic control efficiency and re-routing of electrons into alternative acceptors such as H_2_ production by Hydrogenases or O_2_ photoreduction by FLVs (14, 20, 50). Here, we have further shown that the higher FLVs-mediated PCEF in *pgrl1* mutants channels excess electrons toward O_2_ under N deficiency. PCEF thus counteracts PSI over-reduction, likely facilitating ATP production and at the same time keeping PET active.

### PGRL1/PGR5-controlled CEF contributes to photosynthetic control during N deficiency

The photosynthetic control refers to mechanisms that restrict the PET reactions mostly occurring in response to environmental fluctuations and is typically achieved on the level of the Cyt *b_6_f* upon acidification of the thylakoid lumen (37, 38, 52, 53). The PGRL1-PGR5-controlled CEF increases lumen acidification efficiency and has been shown to contribute to the photosynthetic control in higher plants (26, 28, 54) as well as in microalgae (20–23, 51). The induction of the photosynthetic control results in PSI donor side limitation in the light which is typically determined *in vivo* via P700^+^ optical readouts (55–57). Our results showed higher acceptor side limitation (at the expense of donor side limitation) in *pgrl1* mutants (**Fig. 2 and Supplemental Fig. S4**) indicating that they are affected in the induction of the photosynthetic control. The decrease in Cyt *b_6_f* abundance could be an additional cellular strategy to further restrict the PET toward PSI (higher donor side limitation in wild-type, **Fig. 2 and Supplemental Fig. S4**). While the wild-type successfully relaxes the redox pressure on PSI and degrades Cyt *b_6_f* under N deficiency, *pgrl1* mutants sustained PET and failed to degrade the Cyt *b_6_f*. This dysfunctioning of the photosynthetic control in *pgrl1* (high acceptor side limitation and Cyt *b_6_f* abundance; **Fig. 2 and 4*A***), promotes the recruitment of stromal electron carriers such as FLVs and TAG biosynthesis (in 137AH background). Similar observations were reported under sulphur deprivation where deletion of PGRL1 or PGR5 resulted in sustained linear electron flow toward H_2_ production in *Chlamydomonas* (20, 51). Our work shows that impairing the accumulation of PGRL1 removes a bottleneck of photosynthetic electron flow during N deficiency under non-limiting CO_2_ conditions (**Fig. 1-3; Supplemental Fig. S1-3**). This characteristic of photosynthetic control makes PGRL1/PGR5-controlled CEF a promising target for improving photosynthetic yield under nutrients deficiency. Indeed, the control of photosynthesis under N deficiency has been seen as safety mechanisms protecting cells from phototoxicity (28, 53, 54). Here, we show that thanks to the presence of FLVs, photosynthesis could be improved by removing PGRL1-controlled CEF during N deficiency under non-limiting CO_2_ conditions.

### FLVs become predominant in the absence of PGRL1/PGR5-controlled CEF

So far, a compensation mechanism between PGRL1 and FLVs has been reported during algal adaptation to light or low-CO_2_ conditions (14, 18, 32). Under high light or low-CO_2_ conditions, the increased activity of FLVs in the PGRL1-deficient strain does not improve the net photosynthesis despite its strong efficiency in generating ATP (18, 58). Similar compensation has also been observed in higher plants where orthologous expression of FLVs rescues the *pgr5* mutants phenotype without further improving photosynthesis (54, 59) although some levels of increased biomass were measured in wild-type Arabidopsis expressing FLVs under light fluctuations thanks to the protective role of FLVs in these conditions (60). In contrast, our results suggest that under N deficiency, FLVs and PGRL1 have an antagonistic role. Indeed, the increased activity of FLVs in the PGRL1-deficient strains resulted in higher net photosynthesis (**Fig. 3*D*; Supplemental Fig. S2**), which shows that rather than compensating each other, PGRL1 is indirectly controlling the activity of FLVs by limiting the maximal electron flow capacity. Our observation of the sustained accumulation of Cyt *b_6_f* in the *pgrl1* but not in *pgrl1 flvB* double mutants under N deficiency **(Fig. 4 A and *D*)** could be attributed to the activity of FLVs-mediated PCEF draining electrons from the photosynthetic chain and indirectly preventing Cyt *b_6_f* from degradation. The latter process is catalysed in *pgrl1 flvB* upon PCEF shortage. The presence of strong FLVs activity in *pgrl1* thereby allows maintaining high PET while generating ATP for CO_2_ fixation and reserve formation.

Because CEF and PCEF are dominant pathways for supplying extra ATP in addition to LEF in the chloroplast, we were expecting that the removal of both PGRL1 and FLVs would have more severe consequences on cell physiology and metabolism as it was observed when CO_2_ is limiting (32). Instead, we observed similar photosynthetic activity and TAG biosynthesis in the *pgrl1 flvB* double mutants as their control strains (**Fig. 3 and 5**), i.e. the additional removal of FLVs suppressed the phenotype of PGRL1 deficiency. A third pathway generating the energy required for photosynthetic CO_2_ fixation and ensuring redox dissipation could be operating in the double mutant *pgrl1 flvB*. Both the NDA2-dependent CEF (19, 29) or a chloroplast-mitochondria electron flow (CMEF) (18) could be good candidates. NDA2 protein level was shown to increase during air photoautotrophic N deficiency under atmospheric CO_2_ level and the CEF rate in *Chlamydomonas* was decreased by 50% in *nda2* mutants (29). However, we observed a similar protein level of NDA2 in the *pgrl1 flvB* as their control lines (**Fig. 4*C***). Nevertheless, the protein level might not always correlate with the activity and other regulatory mechanisms (e.g., phosphorylation or redox regulation) can modulate enzyme activity. Additionally, the stimulated mitochondrial respiration rates as well as increased in Cox IIB protein level during N-deficiency in all the strains (except *pgrl1_137AH_*) (**Fig. 4; Supplemental Fig. S3**) points to a strong activity of CMEF (18, 32), which might compensate for CEF and PCEF deficiency in *pgrl1 flvB* N-deprived cells. CMEF may have two distinct roles, either to supply additional ATP or favour redox dissipation. In the context of generating extra ATP, the Cox IIB pathways could take over the AOX1 alternative oxidase pathway because of its efficiency in generating ATP (18). Further investigation would be required to identify the mechanisms generating ATP in the absence of PGRL1 and FLVs. Altogether, we conclude that FLVs maintain high photosynthetic activity under N deficiency in the absence of PGRL1 by channeling excess electrons toward O_2_ meanwhile generating ATP for CO_2_ and downstream metabolic pathways (TAG production).

### Relationship between cellular redox landscape and carbon storage

A major biotechnological challenge in algal domestication for biofuel is the tradeoff between growth and lipid productivity. In *Chlamydomonas* as in many other microalgae, starch and TAG massively accumulate but mostly under stress conditions in particular N deficiency when cell division stops and productivity is impaired. Considerable efforts have focused on the study of the molecular mechanisms behind the onset of reserve accumulation by monitoring omics responses to a stress (5, 61–64), or focused on specific steps of fatty acid and TAG biosynthesis, which have resulted in some limited improvement in productivity (65, 66). Improving productivity requires a better understanding of the crosstalk between photosynthetic carbon fixation, environmental signals and the redox balance, which all govern reserve accumulation (67–69). Here by studying mutants affected in CEF and PCEF, we explored the relationships between the cellular redox status and carbon storage.

The increased accumulation of TAG but not starch observed in the PGRL1-deficient strain (137AH background) under N deficiency (**Fig. 5 *A* and *B***) is a consequence of continuous production of NADPH and ATP through PCEF-dependent PET. In line with our finding, the *Chlamydomonas pgd1* mutants (e.g. *Plastid galactoglycerolipid degradation 1)* with reduced LEF rate (less ATP and NADPH) are shown to produce less TAG under N starvation (11, 70). The report that the *pgrl1* mutant made less TAG than wild-type under mixotrophic conditions (71) is not surprising. It is well known that the bioenergetics of *Chlamydomonas* under photoautotrophic conditions differ from mixotrophic conditions (72–74). The presence of acetate can drastically affect cellular bioenergetics levels i.e. its uptake consumes ATP and its metabolism produces NADH, therefore further favoring oil synthesis (72, 73, 75, 76). TAGs accumulation was different in the CC125 background where the *pgrl1_CC125_* mutant made similar amounts of TAGs as its background strain (**Supplemental Fig. S6*J*)**. This seems not surprising when we consider that TAG accumulation is a metabolic consequence of changes in chloroplast redox state and in photosynthetic performance. The preservation of PSII yield and net O_2_ evolution was less dramatic in the CC125 background than in the 137AH background, which could be one of the reasons behind strain-dependent phenotype. This further reflects the fact that oil content is a complex trait, and it is the consequence of the interaction between genetic makeup, its metabolic flexibility and the capacity of the extent of a given cell’s response to environmental changes.

In contrast to *pgrl1_137AH_*, the *flvB* mutants accumulated lower amounts of TAGs under N deficiency, and no difference in TAG was observed in the *pgrl1 flvB* double mutants, consisting again with the profile in photosynthetic performance and with their antagonistic roles. The *flvB* mutants accumulated higher amount of starch under N sufficient condition though. To conclude, this work further points to the importance of the role redox management on carbon allocation and storage, and that this effect is dependent on many factors including the genetic background, the trophic style, as well as the nutrient status.

## Materials and methods

### Growth conditions and strains

The *pgrl1_137AH_* (from the background 137AH) with its complementing strain (*pgrl1::PGRL1*-2), *flvB* mutants from the background CC-4533 and the *pgrl1 flvB* double mutants were previously described (20, 31, 32). *pgrl1_CC125_* mutants were generated in the CC125 background using CRISPR-Cas9 (see Supplemental Materials and Methods). All the information about the strains used are reported in **Supplemental Table 1**. Cells were routinely cultivated in an incubation shaker (INFORS Multitron pro) maintained at 25°C, with 120 rpm shaking and constant illumination at 50 µmol m^−2^ s^−1^. Fluorescent tubes delivering white light enriched in red wavelength supplied lightings in the INFORS. All experiments were performed under photoautotrophic condition with minimum medium with or without nitrogen source (MM and MM-N) buffered with 20 mM MOPS at pH 7.2 in air enriched with 1% CO_2_. Due to cell aggregation in the 137AH background (notably the *pgrl1* mutant) that prevent accurate cell counting, total cellular volume was measured using a Multisizer 4 Coulter counter (Beckman Coulter) and the different strains were diluted to reach a similar cellular concentration before N deficiency experiments.

### Measurement of chlorophyll fluorescence using a Pulse Amplitude Modulation (PAM)

Chlorophyll fluorescence was measured using a PAM fluorimeter (Dual-PAM 100, Walz GmbH, Effeltrich, Germany) on the MIMS chamber as described in (49) using green actinic light (1250 µmol photon m^-2^ s ^-1^, green LEDs). Red saturating flashes (8,000 µmol photons m^−2^ s ^−1^, 600 ms) were delivered to measure the maximum fluorescence (FM) every 30 s (before and upon actinic light exposure). The maximum PSII quantum yield was calculated as *Fv*/*Fm= (Fm-Fo)/Fm* where F_0_ is the basal fluorescence obtained with the measuring light and Fm the fluorescence emitted after saturating pulse (77). PSII operating yield (ΦPSII) was calculated as ΦPSII= (F_M_’-Fs)/ F_M_’ with F_M_’ the fluorescence value after saturating pulse, Fs the stationary fluorescence during actinic light exposure.

### O_2_ exchange measurement using Membrane Inlet Mass Spectrometry (MIMS)

O_2_ exchanges were measured in the presence of [^18^O]-enriched O_2_ using a water-jacketed, thermoregulated (25°C) reaction vessel coupled to a mass spectrometer (model Prima ΔB; Thermo Electronics) through a membrane inlet system (78). The cell suspension (1.5 mL) was placed in the reaction vessel and bicarbonate (10 mM final concentration) was added to reach a saturating CO_2_ concentration. One hundred microliters of [^18^O]-enriched O_2_ (99% ^18^O_2_ isotope content; Euriso-Top) was bubbled at the top of the suspension just before vessel closure and gas exchange measurements. O_2_ exchanges were measured during a 3-min period in the dark, then the suspension was illuminated at 1250 μmol photons m^−2^ s^−1^ for 5 min using green LEDs followed by 3-min in the dark. Isotopic O_2_ species [^18^O^18^O] (m/e = 36), [^18^O^16^O] (m/e = 34), and [^16^O^16^O] (m/e = 32) were monitored, and O_2_ exchange rates were determined (78). Argon gas was used to correct O_2_ exchange measured by the spectrometer as described in (78).

### Redox state measurements of P700

The PSI primary electron donor, were obtained from optical signals (705 nm – 740 nm) using a Joliot-type spectrophotometer (JTS-150, Spectrologix USA), described in detail elsewhere (23, 79, 80). Harvested cells (2 min, 4000 rpm, 22°C) were resuspended in cuvettes to comparable densities in growth medium containing 20% Ficol (*w*/*v*). Samples were subjected to 2.5-s alterations of dark/light (490 µmol photon m^-2^ s^-1^, 630-nm LEDs), leading to a partial P700 pre-oxidation during the light period (donor side limitation). Additional fractions of photo-oxidizable P700 (PSI yield) were obtained by a 25 ms saturating pulse before the dark period, revealing non-oxidizable P700 (acceptor side limitation) after comparison to fully oxidized P700 when measured in the presence of 20 µM 3-(3,4-dichlorophenyl)-1,1-dimethylurea (DCMU).

All other methods are described in ***SI Materials and Methods*.**

## Acknowledgments

O.D. thanks The French Atomic Energy and Alternative Energy Commission (CEA) for a PhD scholarship. G.P. and Y.L.-B. acknowledge the continuous financial support of CEA (LD-power, CO2Storage). A.B. acknowledges the support from the Carnegie Institution for Science. We thank Bertrand Legeret for maintaining the HelioBiotec lipidomics platform and thank Mallaury Cabanel for assistance in performing immunoblot. We also acknowledge the ZoOM microscopy facility. M.H. (DFG Research Unit FOR 5573) and F.B. (BU 3426/3-1) acknowledge Deutsche Forschungsgemeinschaft for funding.

## Author’s Contributions

Y.L-B., G.P., A.B., and O.D. conceived the study. Y.L-B., G.P. and A.B. provided supervision. O.D. performed most of the experiments. P.A., M.B. and O.D. carried out biochemical experiments. M.B. and O.D. performed starch and lipid analysis. G.P. supervised the MIMS experiments. C.S. and J.I. generated the CRISPR-mediated mutant lines under the supervision of A.B.. F.B. and M.H. performed and analyzed the P700 measurements. O.D. drafted the manuscript with contributions from A.B., G.P, F.B., M.H. and Y.L-B.

