## Supplementary material for "Cyclic and pseudo-cyclic electron pathways play antagonistic roles during nitrogen deficiency in *Chlamydomonas reinhardtii*": SI information: Dao et al. SI information.docx

**ORCID IDs:** 0000-0002-7040-5770 (O.D.), 0000-0001-7434-6416 (A.B.), 0000-0001-6704-6634 (F.B.), 0000-0001-5098-1554 (M.B), 0000‑0002‑3376‑6550 (P.A.), 0000-0002-4078-6250 (C.S.), 0000-0001-9670-6101 (M.H.), 0000-0002-2226-3931 (G.P.), 0000-0003-1064-1816 (Y.L.-B.)

**This PDF file includes:**

Supplemental Materials and methods

Supplemental figures 1 to 6

**Supplemental Materials and methods**

**Generation of *pgrl1* mutant in the CC125 background**

Chlamydomonas mutant showing targeted insertion in the *PGRL1* locus was generated in the CC-125 background (mt+, nit-) using CRISPR/Cas-9 mutagenesis. Single guide RNAs (sgRNAs) (**Supplemental Table 2**) were designed by CHOPCHOP (1) using version 5.6 of the *Chlamydomonas reinhardtii* genome. Ribonucleic proteins (RNPs) were prepared by duplexing the sgRNA (20% volume per volume - v/v-) and Cas-9 (IDT, Ref# 427093062.) (20% v/v) with duplex buffer (IDT, Ref# 325470395) (60% v/v). Prior to electroporation, cells in the CC-125 background were grown in Tris-acetate-phosphate (TAP) at 22 °C under continuous illumination of 50 μmol photons m^−2^s^−1^. After treatment with autolysin for 3 hours, cells were electroporated in the presence of hygromycin resistance cassette (2 µg mL^-1^), and the RNP mixture. The cells were resuspended in 10 mL TAP with 40 mM sucrose and left shaking overnight in dim light (10-20 µmol photons m^-2^s^-1^). Hygromycin resistant transformants were selected on TAP-agar plates containing hygromycin (20 µg mL^-1^). sgRNA targeted regions were amplified via PCR to check for full or partial insertion of the hygromycin resistance cassette at the target site (**Supplemental Table 3 and Fig. 2*D***).

**Starch quantification**

Starch was quantified from 1 mL of culture containing ∼5 million cells. Briefly, cells were harvested by centrifugation at 13000 *g* for 4 min at 4°C. Pellet was resuspended into 1 mL of methanol, mixed vigorously, and stored at −20°C before analysis. The supernatants were removed and the residual methanol was evaporated from the pellets incubated at room temperature in a fume hood. Pellets were resuspended in 400 μL of distilled water and autoclaved for 20 min at 120°C to solubilize the starch polymer. Total starch was quantified using an enzymatic starch assay kit (Sigma-Aldrich; ref. SA-20) following the manufacturer’s instructions. Glucose converted from the starch was quantified using an automated YSI 2700 glucose analyzer (YSI Life Sciences) previously calibrated using a commercial glucose standard.

**TAG quantification**

Exponentially grown cells (~20 millions) were harvested by centrifugation at 4000 *g* for 3 min at 4°C. Pellets were resuspended in 1 mL of hot isopropanol containing 0.01% butylated hydroxytoluene (BHT) (w/v) for 10 min at 85°C to quench lipases (2). A mixture of methyl tert-butyl ether (MTBE):H_2_O (3/1, v/v) was added to extract lipids, vortexed and allow for phase separation. Finally, lipids were recovered from the upper organic phase after centrifugation at 4000 *g* for 3 min at 4°C. Solvent was evaporated under a stream of N_2_ and total lipid extract was dissolved in 200 µL of chloroform:methanol (2:1 v/v) prior to analysis. TAG was separated from other lipid classes by Thin Layer Chromatography (TLC) and quantified based on densitometry method via comparing a standard curve generated from loading known amount of the standard TAG51:0 (17:0/17:0/17:0) (Sigma‐Aldrich) to the same plate as described previously in (3). Total fatty acids were quantified as described in (3). Briefly, part of the extracted lipids was converted to FAMEs by acid-catalysed transmethylation, extracted into hexane and analyzed by gas chromatography coupled to flame ionization detector and mass spectrometry (GC-FID-MS) (Agilent 7890A GC and Agilent 5975C MS, Agilent Technologies, Palo Alto, CA, USA).

**Immunoblot**

For protein extraction, exponentially grown cells (3 mL at around 10 µg mL^-1^ of chlorophyll) were harvested by centrifugation at 4000 *g* for 3 min at 4°C. Pellets were resuspended in 200 µL of 1% SDS, to which 800 µL of cold acetone were added to extract chlorophyll and the suspension was incubated for 30 min at -20°C. Samples were then centrifuged for 10 min, 13000 *g* at 4°C to precipitate proteins. Chlorophyll concentration was measured by spectrophotometry from supernatant. The protein pellet was resuspended with Novex™ Nupage™ LDS buffer 1x (Invitrogen™) containing reducing agent DTT, at a final volume corresponding to 1 µL per 0.1 µg chlorophyll measured (corresponding to 1 µg µL^-1^ of protein), the resuspended proteins were then denatured for 20 min at 70°C. Ten µg of protein were loaded on Novex™ Nupage™ Bis tris 12% (Invitrogen™) gel, migrated 1 h at 190 V in Novex™ Nupage™ MOPS (Invitrogen™) buffer (or Novex™ Nupage™ MES (Invitrogen™) according to protein molecular weight and transferred to nitrocellulose membrane using semidry transfer technique. Immuno-detection was performed using antibodies raised against PGRL1 (4), NDA2 (5). Other antibodies were ordered from Agriserea for PSBD (AS06 146) 1/10000, PSAD (AS09 461), Cyt *f* (AS06 119) 1/500, Cox IIB (AS06 151) 1/5000, ATG8 (AS14 2769) 1/2000 (6, 7), 50S ribosomal protein L30 (AS08 331) 1/1000 and α-Tubulin (AS10 680) 1/1000. Secondary anti-rabbit peroxidase-conjugated antibodies (Sigma-Aldrich; no. AQ132P) (1/10,000) were used for the detection with the G:BOX Chemi XRQ system (Syngene) using ECL detection reagents (GE Healthcare). Images were captured with a CCD camera equipped with a GeneSys Image Acquisition Software (Syngene).

**Statistical analysis**

One-way ANOVA using GraphPad Prism (GraphPad Software) was used to perform statistical analysis. The *p* values were computed by One-way ANOVA test (uncorrected *p* values) or using Turkey corrections for large datasets (adjusted *p* values). Shown are grouping of strains into statistical families according to *p* values indicated by asterisk (* for *p* ≤ 0.05; ** for *p* ≤ 0.01; *** for *p* ≤ 0.001 and **** for *p* ≤ 0.0001) or letters indicating parameter-specific significances. The statistical significance (α) was set as 0.05 (5%).

**Supplemental Tables**

**Table S1: List and origins of mutants used in this study**

| **Gene ID** | **Strains** | **Genetic background** | **Mutant generation** | **Reference article** |
| --- | --- | --- | --- | --- |
| **Cre07.g340200**  **(PGRL1)** | *pgrl1_137AH_* | 137AH | Random insertion | (4) |
|  | *pgrl1_CC125_* | CC125 | CRISPR-Cas9 | This article |
| **Cre16.g691800**  **(FLVB)** | *flvB-21* | CC4533 | Random insertion (CLiP) | (8) |
|  | *flvB-208* | CC4533 | Random insertion (CLiP) | (8) |
|  | *flvB-308* | CC4533 | Random insertion (CLiP) | (8) |
| **Cre07.g340200/Cre16.g691800**  **(PGRL1 and FLVB)** | WT1 | 137AH x CC4533 | Mating between *pgrl1_137AH_* and *flvB-21* | (9) |
|  | WT3 | 137AH x CC4533 | Mating between *pgrl1_137AH_* and *flvB-21* | (9) |
|  | *pgrl1 flvB-3* | 137AH x CC4533 | Mating between *pgrl1_137AH_* and *flvB-21* | (9) |
|  | *pgrl1 flvB-5* | 137AH x CC4533 | Mating between *pgrl1_137AH_* and *flvB-21* | (9) |

**Table S2: sgRNAs to knockout target genes**

| **Mutant** | **Gene ID** | **sgRNA Sequence** | **Target Exon** |
| --- | --- | --- | --- |
| ***pgrl1_CC125_*** | Cre07.g340200 | CTTGCCCTCGTAGTACCACGAGG | 1 |

**Table S3: Primer pair sequences for genotyping mutants**

| **Target Gene** | **Target Exon** | **Primer Name** | **Oligo Sequences** | **Predicted Band Size in CC-125 (bp)** |
| --- | --- | --- | --- | --- |
| ***PGRL1*** | 1 | oAB0013 | GCAATGTTGCTGCTGACTACTG | 225 |
|  | 1 | oAB0014 | AGTCCTCGTCAGTCATAATGGG |  |
| ***STT7*** | 1 | oAB0033 | GCACGAACCAAGACACACATAG | 237 |
|  | 1 | oAB0034 | GTAGACGATGTCACCGCACTT |  |

**Supplemental Figures**

**Supplemental Figure 1.** **PSII activity measurement using PAM in *pgrl1_137AH_*, *flvB* and *pgrl1 flvB* mutants upon N deficiency**

**A,** Generation of *pgrl1 flvB* double mutants by crossing and their genealogy. Antibodies against PGRL1, FLVA and FLVB were used to confirm the genotype. RbcL is used as a positive control.

**B-C**, Maximum quantum efficiency and the operating yield of PSII were measured from 15min dark-adapted cells of *pgrl1_137AH_* and 137AH after 0, 22, 28, 46 and 50 h of N deficiency and 12 h after N resupply. Shown are an average of at least three biological replicates ± SD.

**D-E**, Maximum quantum efficiency and the operating yield of PSII were measured from 15min dark-adapted cells after 48 h of N deficiency. PSII yield were calculated after 4min of illumination using green actinic light (1250 µmol photon m^-2^ s ^-1^, green LEDs). Shown are an average of at least three biological replicates ± SD. Asterisks represent statistically significant difference comparing mutants with their control strains (* *p*<0.05, ** *p*<0.01 and *** *p*<0.001) using one-way ANOVA.

**Supplemental Figure 2.** **O_2_ exchange measurement under N replete and deficiency in *pgrl1_137AH_*, *flvB* and *pgrl1 flvB* mutants**

**A**, O_2_ exchange under N replete.

**B,** O_2_ exchange under N deficiency.

O_2_ exchange rates were measured during the dark-light-dark transition from N replete and deficient cells of *flvB*, *pgrl1 flvB* and respective control lines using a MIMS in the presence of [^18^O]-enriched O_2_. Net O_2_ evolution (green) was calculated as gross O_2_ evolution (blue) – gross O_2_ uptake (red). Shown are an average of at least three biological replicates ± SD.

**Supplemental Figure 3.** **Photosynthetic activity in CRISPR-Cas9 generated *pgrl1_CC125_* mutant**

**A-B** Chlorophyll fluorescence during the dark-light-dark transition under N replete and after 16h of N deficiency.

**C**, PSII operating yield N replete and after 16h of N deficiency.

**D-F**, O_2_ exchange rates were measured using a MIMS during the dark-light-dark transition from N-replete (**D**) and N-deficient (**E**) conditions. Net O_2_ evolution (green) was calculated as gross O_2_ evolution (blue) – gross O_2_ uptake (red).

**G**, Genetic characterization of mutants using PCR on genomic DNA. Primers and expected size are listed in **Supplemental Table 1 and 2**. STT7 was used as a control.

**H**, Immunoblot showing the accumulation of PGRL1, Cyt *f*, PsaD, Nda2 and Tub in *pgrl1_CC125_* under N replete and after 16h of N deficiency.

Wild type CC125, *pgrl1_CC125_* N-replete cells cultivated photoautotrophically with 1% CO_2_ in air under continuous light (50 μmol photons m^−2^ s^−1^) were transferred into N-free media for 16h prior to measurements. PSII yield were calculated from (**A** and **B)** after 4min of illumination using green actinic light (1250 µmol photon m^-2^ s ^-1^, green LEDs). Shown are an average of at least three biological replicates ± SD. Asterisks represent statistically significant difference comparing mutants with their control strains (* *p*<0.05, ** *p*<0.01 and *** *p*<0.001) using one-way ANOVA.

**Supplemental Figure 4.** **Redox state measurements of P700 in *pgrl1_CC125_* mutant**

The influence of PGRL1 on the redox state of the primary PSI donor, P700, was monitored in the genetic background CC125 **(A-B)**. Panel A shows raw kinetics (red/yellow/black bars: 490/3000/0 μmol photons m^−2^ s^−1^) in the presence and absence of PSII inhibitor DCMU (means ± SD of 4 biological replicates). Panel B quantifies the corresponding P700 pools at the end of the 2.5-s light period (letters indicate parameter-specific significances using one-way ANOVA/Fisher-LSD, *p*<0.05). At the time of quantification, P700 remained photo-oxidizable by a saturating pulse (PSI yield), was redox inactive (no acceptors) or was pre-oxidized (no donors). Wild-type CC125, *pgrl1_CC125_* N-replete cells cultivated photoautotrophically with 1% CO_2_ in air under continuous light (50 μmol photons m^−2^ s^−1^) were transferred into N-free media for 24 h prior to measurements.

**Supplemental Figure 5.** **Evaluation of biomass and carbon storage during N deficiency in *pgrl1_137AH_***

**A**, Kinetics of starch accumulation during N deficiency.

**B**, Kinetics of TAG accumulation during N deficiency.

**C**, Cell growth as a function of cellular volume during N deficiency.

**D**, representative images of confocal microscopy observation of lipid droplet stained with Bodipy dye. Pseudo-colors were used: BODIPY-stained LDs in yellow, chlorophyll autofluorescence in red. DIC: differential interference contrast. Scale bar = 5 µm.

Cells, cultivated photoautotrophically with 1% CO_2_ in air under continuous light of 50 μmol photons m^−2^ s^−1^, were transferred into N-free media and samples were harvested at different times as indicated on the graphs. Shown are an average of at least three biological replicates ± SD. Asterisks represent statistically significant difference compared to the control 137AH (* *p*<0.05, ** *p*<0.01 and *** *p*<0.001) using one-way ANOVA.

**Supplemental Figure 6.** **Starch and TAG production in *pgrl1_137AH_, flvB, pgrl1 flvB* and *pgrl1_CC125_***

**A-C** Starch quantification in N replete cells of *pgrl1_137AH_*, *flvB* and *pgrl1 flvB* mutants.

**D-F** TAG quantification in N replete cells of *pgrl1_137AH_*, *flvB* and *pgrl1 flvB* mutants.

**G-J** Starch and TAG quantification in N replete and after 24 hours of N deficiency in *pgrl1_CC125_*.

Cells were cultivated photoautotrophically with 1% CO_2_ in air under continuous light of 50 μmol photons m^−2^ s^−1^. Shown are an average of at least three biological replicates ± SD. Asterisks represent statistically significant difference comparing mutants with their control strains (* *p*<0.05, ** *p*<0.01 and *** *p*<0.001) using one-way ANOVA.
