## Supplementary figures and images for "Cyclic and pseudo-cyclic electron pathways play antagonistic roles during nitrogen deficiency in *Chlamydomonas reinhardtii*"

### Supplemental Fig. 1.tif

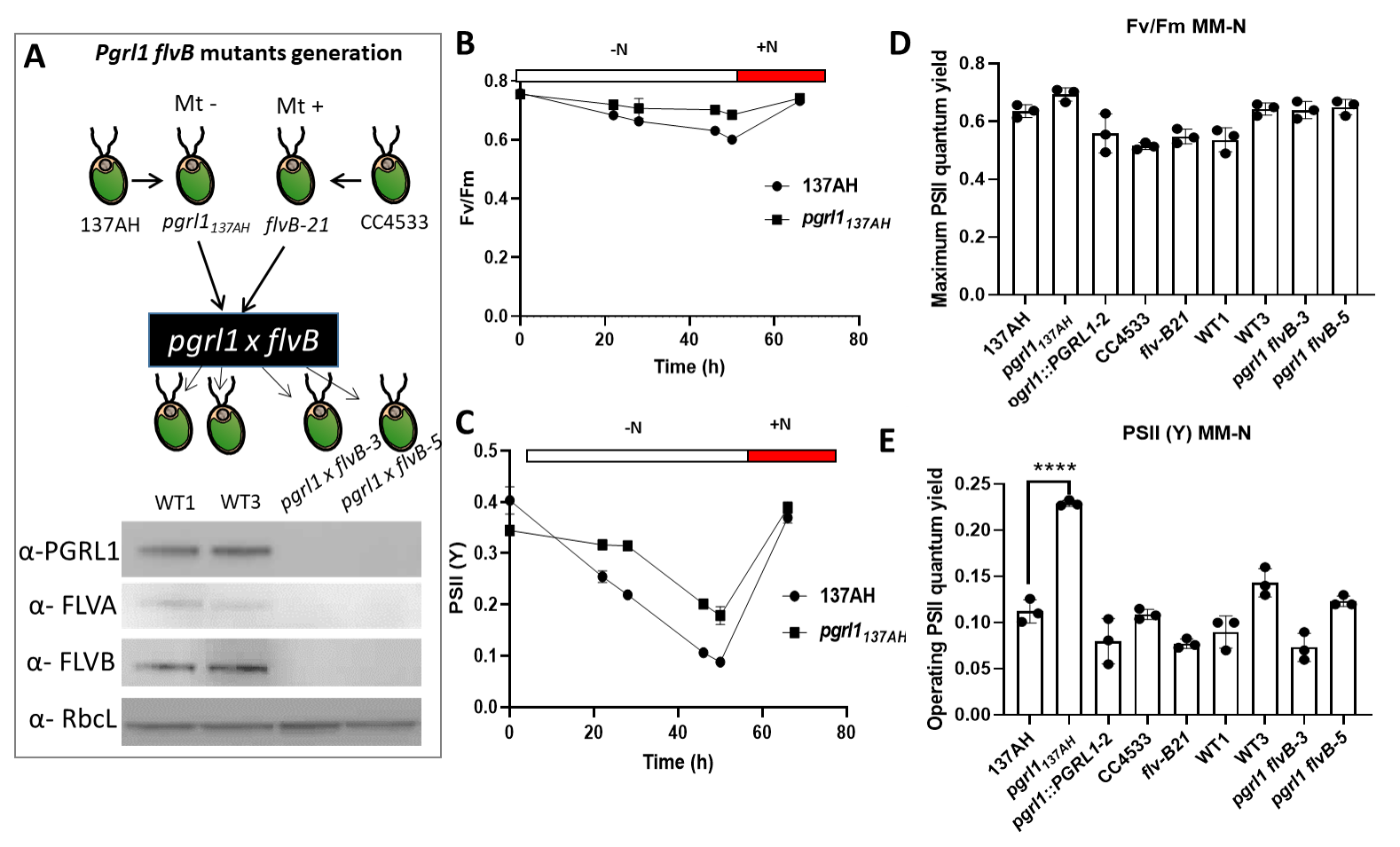

### Supplemental Fig. 2.tif

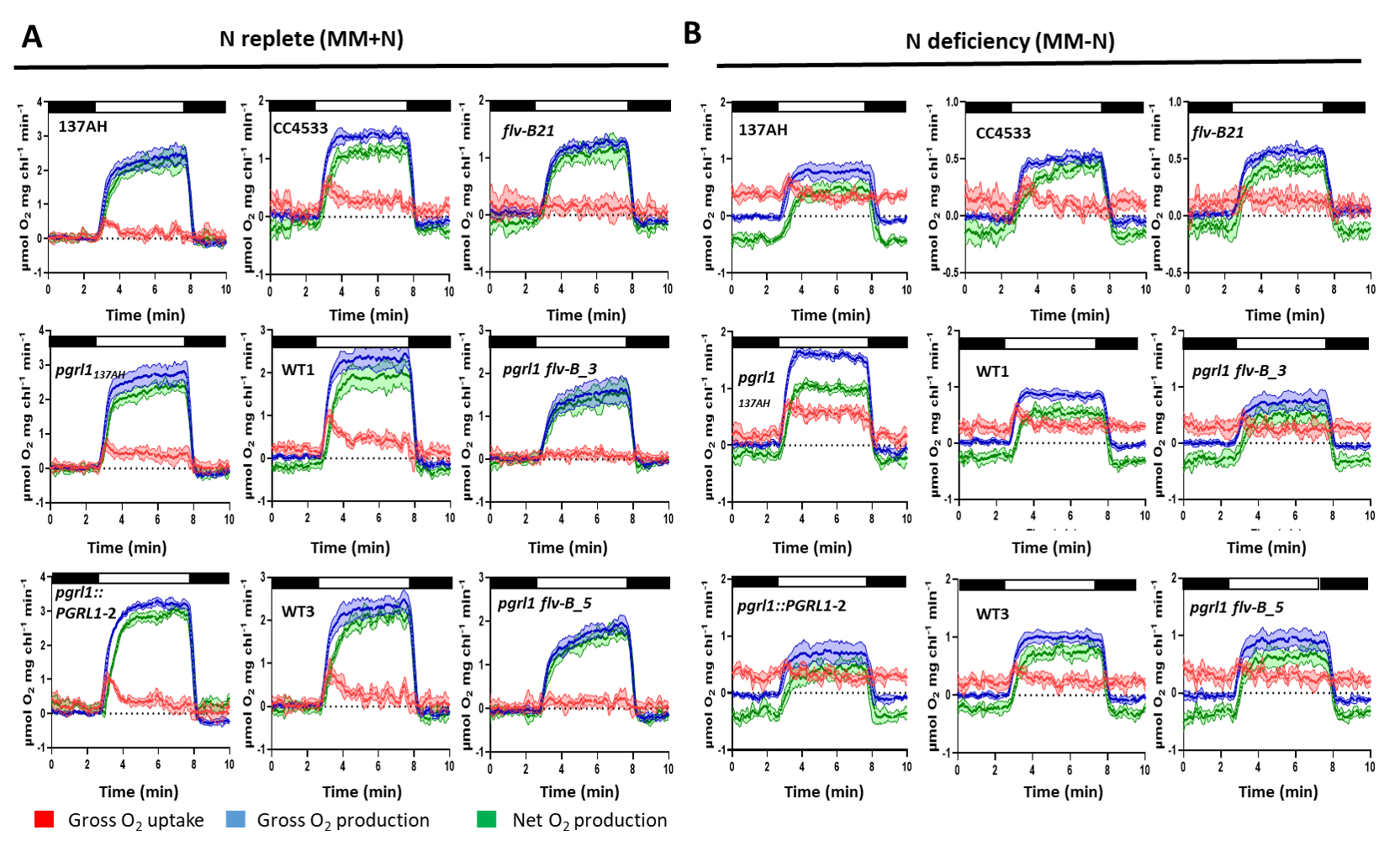

### Supplemental Fig. 3.tif

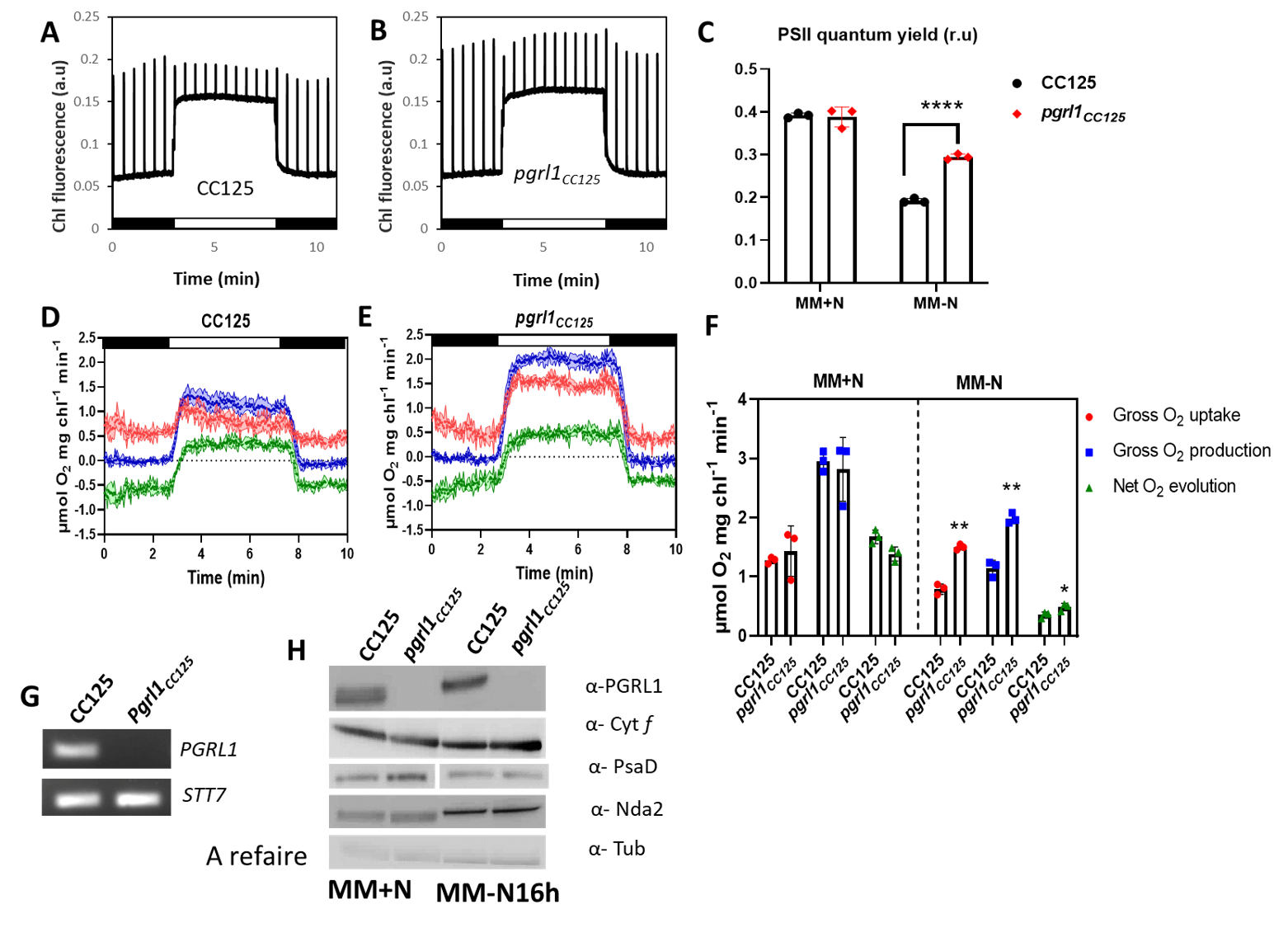

### Supplemental Fig. 4.tif

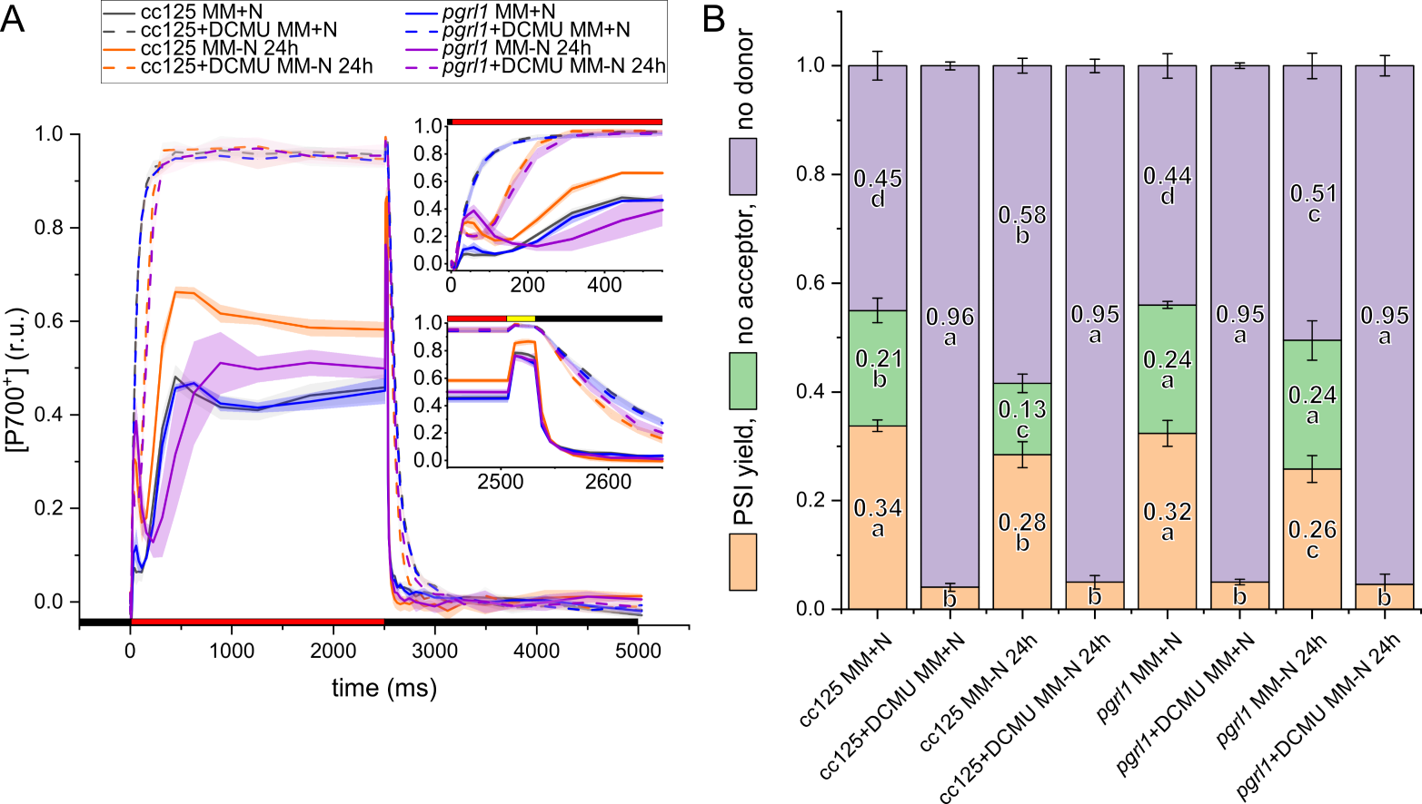

### Supplemental Fig. 5.tif

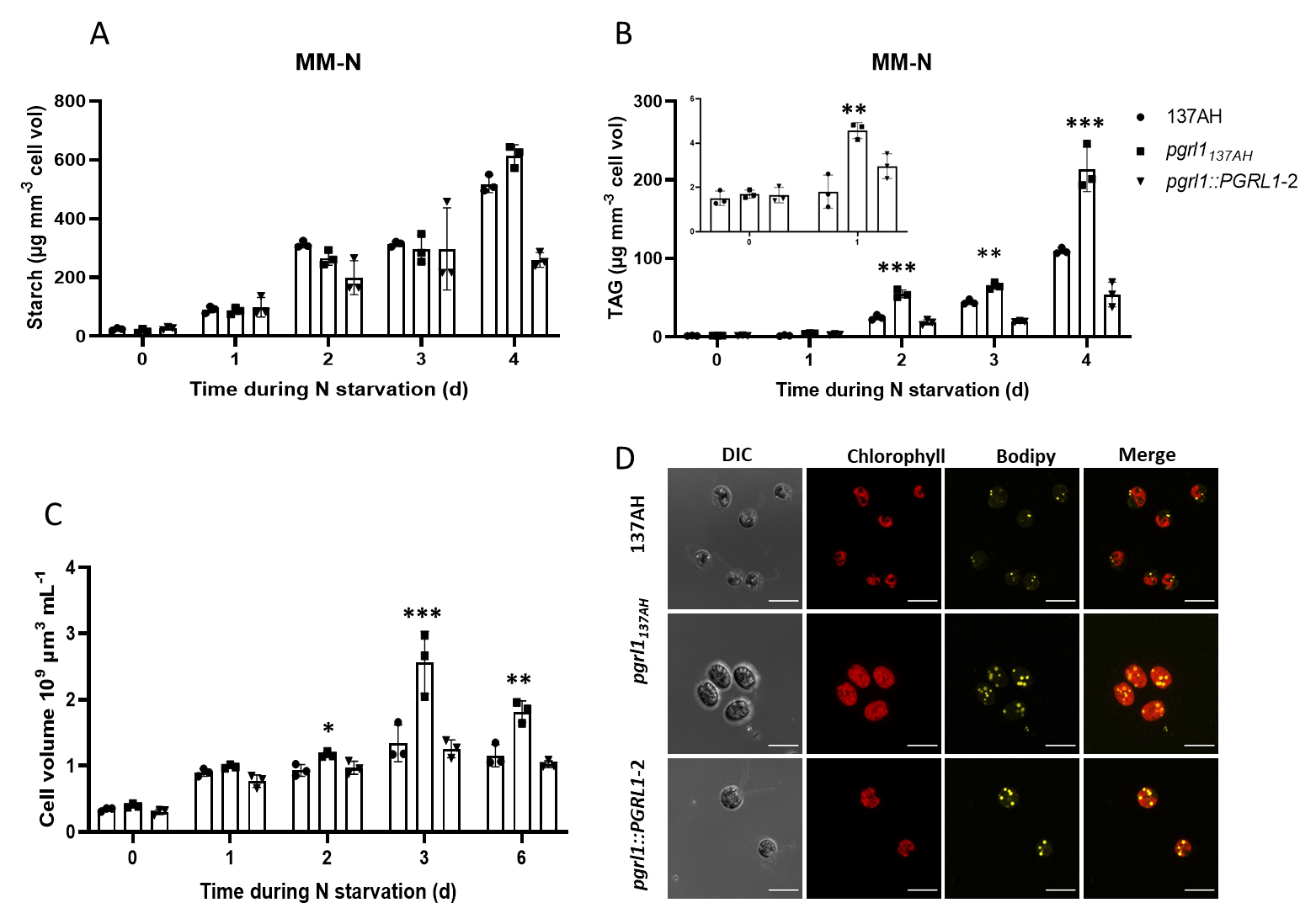

### Supplemental Fig. 6.tif

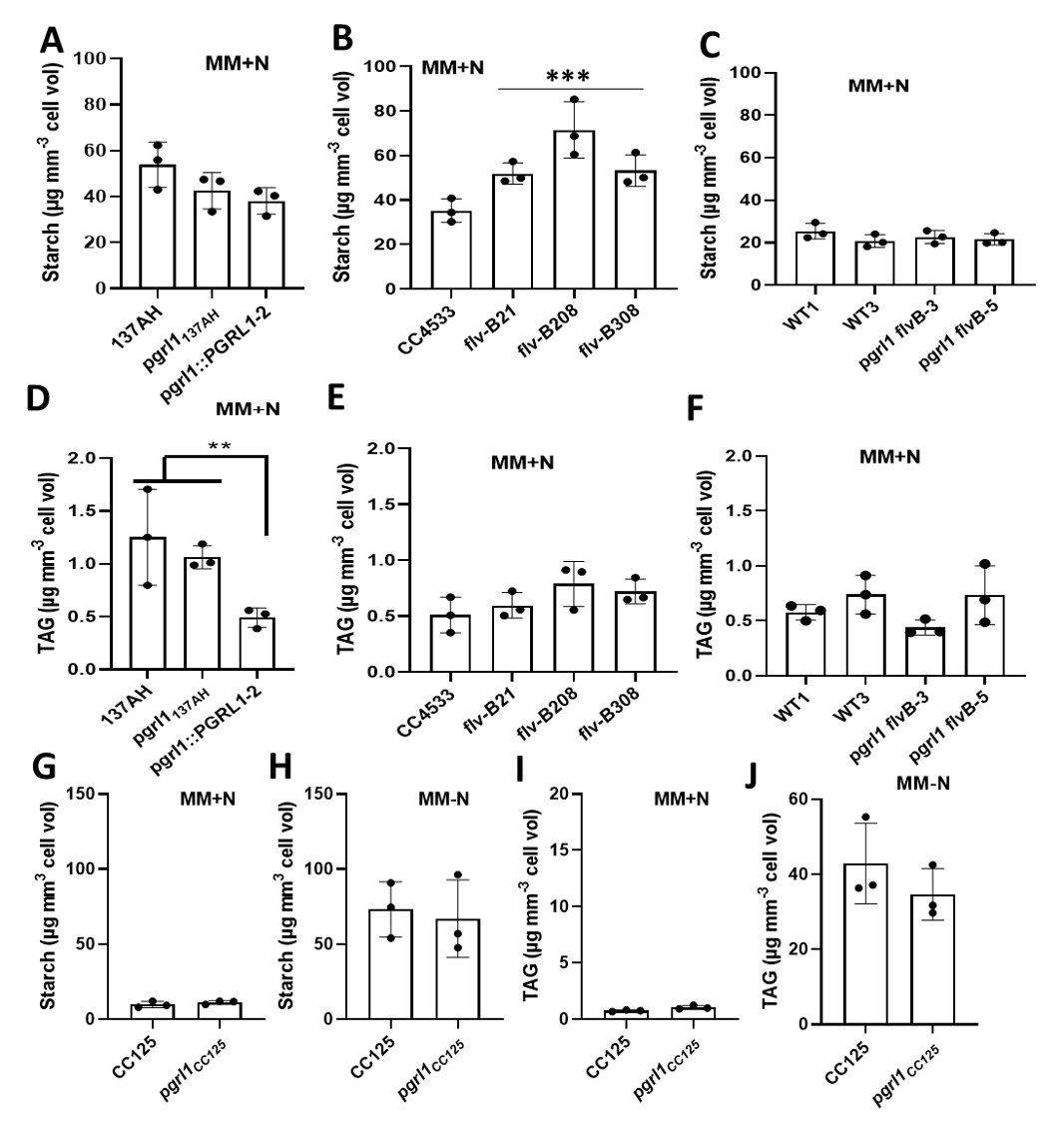
